# Functional validation of the *Plasmodium falciparum* K13 C580Y mutation in recently collected Ethiopian isolates

**DOI:** 10.64898/2026.03.17.712112

**Authors:** Angana Mukherjee, Ashenafi Bahita Assefa, Christopher V Turlo, Lisa Checkley Needham, Douglas Shoue, Tarrick Qahash, Mahlet Belachew, Dagimawie Tadesse, Enirsie Kassie, Mulu Berihun, Bokretsion G. Brhane, Jonathan B. Parr, Michael T. Ferdig, members of the MAREE Consortium, Sisay Adane, Adisu Tesfaye, Jenna Zuromski, Abebe A. Fola, Geremew Tasew, Getachew Tollera, Jonathan J. Juliano, Jeffrey A. Bailey

**Author notes:** These authors contributed equally (Angana Mukherjee, Ashenafi Bahita Assefa). Corresponding author: Angana Mukherjee.

## Abstract

Recent genomic investigation in Ethiopia identified the first detection of the *Plasmodium falciparum* kelch13 (K13) C580Y substitution in the Horn of Africa. To assess its functional impact, we introduced C580Y into two recently collected Ethiopian clinical isolates using CRISPR-Cas9 genome editing. Ring-stage survival assays showed significantly elevated in vitro dihydroartemisinin (DHA) survival in edited parasites relative to isogenic controls, demonstrating that C580Y confers artemisinin tolerance in contemporary Ethiopian parasite genetic backgrounds.

## Text

A recent genomic investigation in northwestern Ethiopia reported the first detection of the *Plasmodium falciparum* Kelch13 (K13) C580Y substitution in the Horn of Africa, a validated molecular marker of artemisinin partial resistance (ART-R). Following its emergence in western Cambodia in the late 2000s, K13 C580Y rapidly spread (1, 2) across the Greater Mekong Subregion (GMS), reaching near-fixation in several countries and displacing other resistance-associated variants. Its repeated association with robust ART-R and efficient spread in low-transmission settings under dihydroartemisinin (DHA)–piperaquine selection highlights its evolutionary success (3, 4). For a K13 substitution to become prevalent in a parasite population, it must satisfy two minimal criteria: it must confer sufficient ART-R to provide a selective advantage under drug pressure, and it must remain compatible with asexual growth and transmission fitness. The evolutionary success of the K13 C580Y variant in Southeast Asia may reflect this balance.

Despite the widespread emergence of multiple K13 substitutions in Africa (5–12), C580Y has only been detected sporadically. However, Zeleke *et al*. recently identified two C580Y mutant infections from the Gondar Zuria district in northwestern Ethiopia, collected between November 2022 and October 2023, and confirmed by molecular inversion probe sequencing and long-read whole-genome sequencing. Haplotype analysis indicated a distinct genetic background relative to the Southeast Asian C580Y lineages, marking the first detection of this mutation in the Horn of Africa to our knowledge (13).

This rare detection raises an immediate biological question: whether C580Y can resist artemether-lumefantrine pressure in patients, establish and persist in African parasite populations or is transient? Addressing this question requires functional evaluation in relevant parasite genetic backgrounds, as the ART-R phenotype associated with K13 substitutions is strongly context dependent. For example,CRISPR-Cas9 introduction of the prevalent Ugandan K13 substitutions A675V and C469Y into both Southeast Asian (Dd2) and East African (MAS-136) parasite lines showed that these substitutions did not uniformly confer reduced DHA susceptibility, particularly in the African background (14). These findings indicate that such K13 substitutions alone may be insufficient to generate resistance, and that permissive genetic backgrounds or additional genetic modifiers are required for phenotypic changes. The introduction of K13 C580Y produced variable phenotypes in long-adapted isolates, conferring resistance in Cambodian, Dd2 (15), Vietnamese (V1/S) (16), Ugandan isolates, and 3D7 (17), but failed to do so in others, such as Tanzanian (F32) (17) and Chinese (FCC1/HN) (18) isolates. Collectively, these studies demonstrated that the ART-R phenotypic consequences of K13 substitutions cannot be inferred from their presence alone; they must be evaluated in the context of relevant genetic backgrounds from the same region or country.

We investigated the impact of K13 C580Y by introducing it into two recently collected clinical isolates of Ethiopian *P. falciparum* and assessed ART-R *in vitro*. The isolates, AM001 (Arba Minch, South Ethiopia Regional State) and SK2 (Selekleka, Tigray, northern Ethiopia), were collected in 2024. Venous blood (5-7 mL) from *P. falciparum* infected patients was collected in sterile heparinized tubes, washed with incomplete RPMI medium, and cryopreserved in glycerolyte within 24 hours of collection as per the WWARN guidelines. The isolates were subsequently culture-adapted *in vitro* and confirmed to carry wild-type K13 propeller sequences using Sanger sequencing (**Supplementary Figure 1**). Using CRISPR-Cas9 genome editing, we edited *k13* for C580Y in both the isolates. As editing controls, we also introduced two synonymous shield mutations in *k13* that did not alter the amino acid sequences (19). Editing of *k13* was performed using the pDC2-coSpCas9-k13guide-gRNA-h*dhfr* all-in-one plasmid containing a *P. falciparum* codon-optimized Cas9 sequence, a human dihydrofolate reductase (h*dhfr*) gene expression cassette (conferring resistance to WR99210) and donor template (containing the respective nonsynonymous and shield/binding site mutations or only shield/binding site mutations) (20). 80 µL of packed RBCs containing 5% ring parasitemia were electroporated with 50 µg of the purified plasmid resuspended in Cytomix using a BioRad Gene Pulser Xcell Electroporation System. Transfected parasites were maintained under 2.5 nM WR99210 (Jacobus Pharmaceuticals) to select edited parasites until 6 days post-transfection, after which the drug was removed from the media. Parasite cultures were microscopically monitored for recrudescence for up to six weeks post-electroporation.

Successful editing was confirmed using Sanger sequencing (**Supplementary Figure 1**). To our knowledge, this is one of the first functional validations of a K13 substitution performed through genome editing in recently collected African patient isolates. By assessing native Ethiopian parasite genetic backgrounds, this approach directly tests whether C580Y confers ART-R in epidemiologically relevant lineages and provides a critical functional context for interpreting the emergence of this mutation reported by Zeleke *et al*. A DHA survival screen performed at 700 nM DHA with a qPCR readout at 120 hours post-exposure (21) demonstrated significantly elevated survival in C580Y-edited lines relative to their isogenic controls (SK2: *p* = 0.0445; AM001: *p* = 0.0031, unpaired Welch’s *t*-test), indicating that C580Y is sufficient to confer artemisinin tolerance in contemporary Ethiopian parasite backgrounds (**Figure 1**).

**Figure 1:**
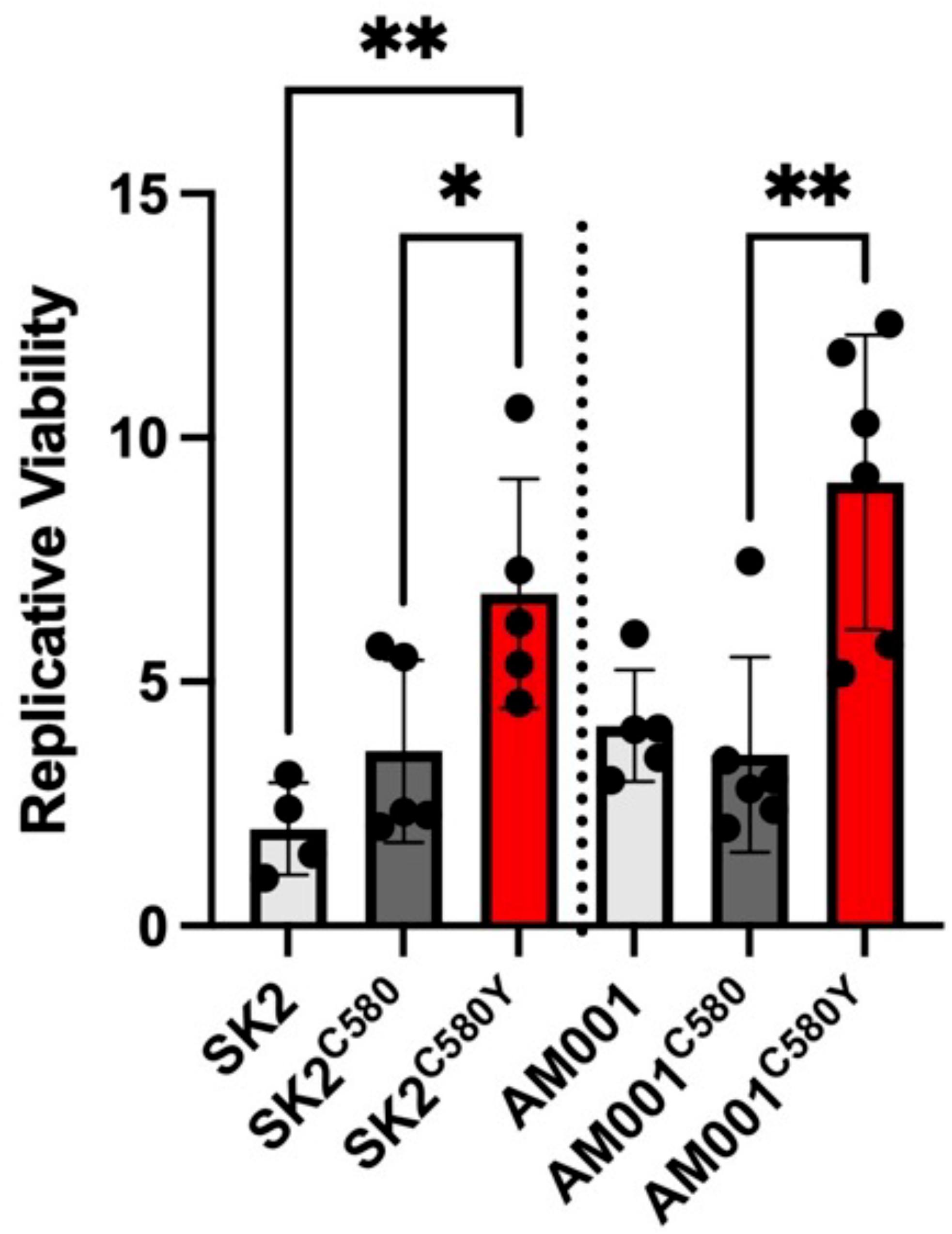
Parasite Survival after DHA exposure in CRISPR-Cas9 C580Y *k13* edited Ethiopian isolates. Mean Replicative Viability from a qPCR read out 120h post-exposure to 700 nM DHA among edited parasites in 4-6 biological replicates is plotted. Error bars indicate the standard error of mean. RBCs infected with Percoll synchronized mature schizonts were incubated with uninfected RBCs for 4-6 hours, after which parasitemia was assessed by flow cytometry. 0.2% parasitemia in 2% hematocrit was exposed to 700 nM DHA or dimethylsulfoxide (DMSO, vehicle control) for 6 hours. The drug was removed by washing, and cultures were maintained for 120 hours. The numbers of intraerythrocytic parasites were measured by SYBR green in a qPCR format and replicative viability after drug treatment was calculated relative to DMSO-treated parasites. Unpaired t-tests with Welch’s correction were used to compare edited *k13* mutant strains with silent edited isogenic strains and their corresponding parental strains (SK2, a patient *P. falciparum* isolate collected from Selekleka Health Center in Tigray and AM001 from Arba Minch in South Ethiopia Regional State).

Together, these findings raise concern that C580Y in Ethiopia could contribute to clinically relevant ART-R, as previously observed in Southeast Asia. Determining the asexual growth and transmissible fitness of these isogenic C580Y-edited lines will be essential to assess whether this newly emergent mutation is likely to persist locally or expand. Such studies will be critical for distinguishing transient emergence from evolutionary success and for anticipating the public-health significance of C580Y in Africa.

## Funding and Acknowledgements

This study was supported by the National Institutes of Health, NIAID (R01AI177791 to JBP). We thank Dr. Marcus Lee for providing the CRISPR plasmids. We also thank Boja Taddese for sample collection and cryopreservation at EPHI.

## Author Contributions

AM contributed to the *in vitro* culture adaptation, transfection, genome editing, sequencing, visual representation of the data, and drafting and editing of the manuscript. LCN contributed to the *in vitro* culture adaptation, CVT, TQ and LCN to the RSA of isolates and edited lines. CVT contributed to the production of DNA for transfections. DS supervised and analyzed the RSA data. JJJ and JAB contributed to the editing of the report. MBelachew, DT, EK, M Berihun, BGB and AA coordinated field work, collected patient isolates and cryopreserved them in glycerolyte. JBP supervised coordinated, reviewed and edited the report. AM and MTF supervised t laboratory experiments. Listed members of the MAREE consortium reviewed the manuscript and contributed to field work, laboratory analyses, and/or laboratory capacity building pertinent to this work.

## Ethics Statement

The study was conducted using malaria parasite samples collected in Ethiopia. Written informed consent was obtained from the participants or their legal guardians prior to sample collection, and the samples were deidentified. The protocol adhered to the principles of the Declaration of Helsinki and relevant national ethical guidelines. The protocol was approved by EPHI’s ethical review board (EPHI-IRB-535-2023).

