## Supplementary Figure 1 for "Functional validation of the *Plasmodium falciparum* K13 C580Y mutation in recently collected Ethiopian isolates"

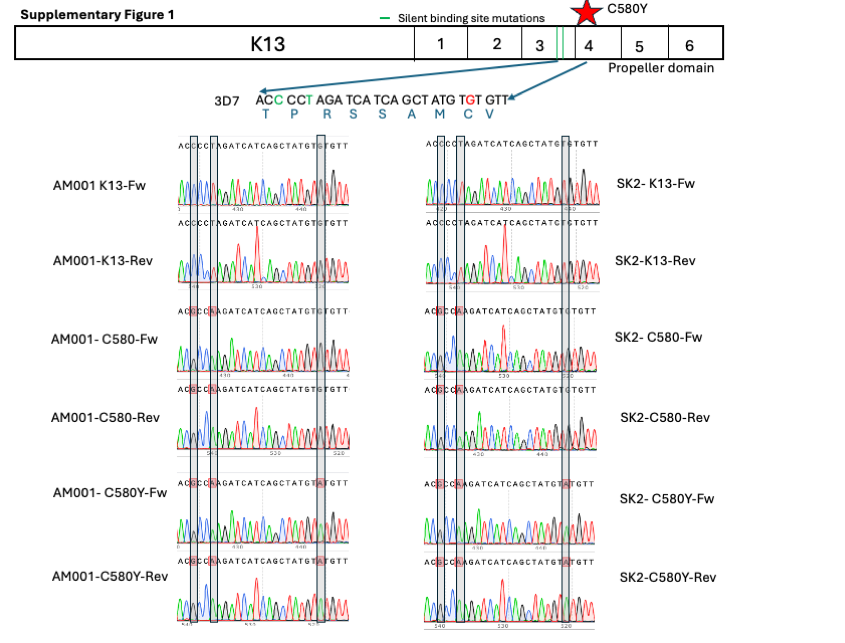

**Supplementary Figure 1:** **CRISPR-Cas9 genetic editing of K13 C580Y and silent binding site control mutations in AM001 and SK2 parasites**. The top schematic shows the locations of the C580Y and control substitutions in the K13-propeller domain. The reference DNA sequence of 3D7 is shown at the top, and the Sanger sequence and chromatogram analysis of forward and reverse alleles of parent and edited lines are shown below. The highlighted bases show the WT and edited alleles.
